# Transcriptional reprogramming of tumor-infiltrating T cells during PD-1 blockade revealed through gene regulatory network and trajectory inference in squamous cell carcinoma

**DOI:** 10.1101/2025.10.15.682506

**Authors:** Roger Casals-Franch, Jordi Villà-Freixa, Lara Nonell, Adrián López García de Lomana

## Abstract

Understanding the tumor microenvironment is crucial for optimizing anti-cancer immune responses. At single-cell resolution, trajectory inference methods can reconstruct the dynamic transitions between cell states during differentiation. Immune checkpoint blockade (ICB) therapies, such as PD-1/PD-L1 inhibitors, are used across multiple cancers, including non-melanoma skin cancers (NMSCs), yet the transcriptional mechanisms that shape T cell responses in this context remain unclear. Here, we analyzed a publicly available squamous cell carcinoma (SCC) single-cell RNA-seq dataset comprising 25,581 tumor-infiltrating T-cell profiles to map differentiation trajectories before and after anti-PD-1 therapy. In CD8^+^ T cells, therapy enhanced the transition from memory to activated states, prominently involving IL-12–associated pathways, and revealed a distinct memory-to-exhaustion trajectory driven by EOMES and TCF7 regulatory activity. Gene regulatory network inference further revealed therapy-induced transcriptional rewiring distinguishing precursor exhausted (Tpex) from terminally exhausted (Tex) states. CD4^+^ T cell populations also underwent substantial reshaping, with trajectory and functional analyses highlighting therapy-driven programs that enhanced CXCL13^+^ Tfh responses while generating fewer but more transcriptionally active Tregs. Together, these findings reveal a dual remodeling of helper and cytotoxic T cell compartments upon PD-1 blockade, define key transcriptional regulators controlling cell-state transitions, and identify potential molecular targets and biomarkers to predict and enhance treatment response.

## Introduction

Cancer immunotherapy (IT) is a central strategy for treating cancer by modulating the immune system response to tumor cells [1]. Among the numerous IT options available, immune checkpoint inhibitors (ICIs) have shown significant clinical effectiveness in treating various types of cancer [2]. However, ICI treatment provides lasting benefits only in a minority of cancer patients while posing a risk of substantial, potentially life-threatening, autoimmune side effects [3]. Thus, there is a pressing need to deepen our understanding of the molecular mechanisms driving the tumor microenvironment (TME) for more efficient immune responses against cancer [4].

Squamous cell carcinoma (SCC) is a type of cutaneous malignancy that represents the leading cause of metastatic disease and mortality among non-melanoma skin cancers (NMSC). While surgical excision, radiotherapy and chemotherapy are the standard-of-care approach, the emergence of IT is improving therapeutic strategies [5].

The effectiveness of immunotherapy depends on the composition of tumor-infiltrating T cells, which undergo complex transcriptomic, epigenomic, and clonotypic changes [6]. CD8^+^ effector T cells in the tumor microenvironment express the PD-1 receptor, and its interaction with PD-L1 on tumor cells leads to loss of function. Blocking the PD-1 pathway prevents CD8^+^ T-cell inhibition, therefore enabling antigen presenting cells (APC) or tumor cells to present antigens through major histocompatibility complex (MHC) molecules, resulting in T cell expansion and ultimately tumor cell killing [7]. Yet both CD4^+^ and CD8^+^ T cells play crucial roles against tumors. CD4^+^ T cells orchestrate immune responses by producing cytokines with immunoprotective properties, while CD8^+^ T cells exert cytotoxic effects to eliminate infected or tumor cells [8]. Chronic PD-1 signaling and elevated PRDM1 expression in CD8^+^ T cells suppress TCF7, promoting T cell exhaustion—a dysfunctional state marked by reduced cytotoxic activity and impaired immune response [9,10]. This phenomenon is characterized by reduced T cell activity and contributes to cancer progression. ICIs therapies aim to reverse T-cell dysfunction and restore effective cytotoxic activity against tumor cells [11]. Recent research has identified a specialized T cell subtype, known as precursor exhausted cells (Tpex), which exhibits features of both exhausted and memory cells and can differentiate into terminally exhausted cells [12]. Tpex cells are crucial for the response to ICIs therapies, with an increased presence of Tpex cells being associated with better patient survival outcomes [12].

The diversity, heterogeneity, and complex interactions in the tumor microenvironment (TME) pose major challenges for characterizing cell-state dynamics and identifying the regulatory mechanisms driving ICI outcomes. Trajectory inference and pseudotime analysis have emerged as a popular approach for reconstructing cell differentiation dynamic trajectories from transcriptome profiles at single-cell resolution [13,14] and has been applied across diverse biological systems from human organoids [15] to cancer immunotherapies [16]. Pseudotime can be considered as a time-like latent variable that indicates the progression of a cell state along the differentiation process [17]. Assuming sufficient coverage over various cell states (e.g., naive, intermediate, memory and mature T cells) this approach has the potential to reveal cell fate decisions of distinct cell subsets, facilitating the discovery of the molecular mechanisms regulating these complex transitions [18–22].

Cell state transitions, including immune cell responses such as effector functions, emerge from complex regulatory interactions across large gene collections [23]. Gene regulatory network (GRN) inference algorithms are computational tools that address such complexity by identifying coherent gene modules and predicting regulatory mechanisms [24]. These tools have demonstrated to be relevant to a variety of clinical challenges such as SARS-CoV-2 infection [25], drug repurposing [26] and cancer resistance and therapy escape [27].

A full understanding of the molecular regulatory mechanisms and interactions governing T cell state transitions in the context of immunotherapies remains largely unknown. In this direction, trajectory inference analysis is a useful framework that has delivered relevant insights. For instance, Yan et al. [28] inferred cell trajectories to characterize the dynamics of transcriptional regulators in T cell exhaustion upon ICI therapy. Through the analysis of gene expression across CD8^+^ T cell transitions into dysfunction, their study revealed a bifurcation from naive to dysfunctional and effector states and identified transcription factors (TFs) associated with these transition stages. Notably, known TFs associated with Tpex such as TCF7 and EOMES were identified, along with TSC22D3, which is associated with improved survival outcomes at patient level. However, understanding both CD4^+^ and CD8^+^ T cell dynamics is essential for a comprehensive view of T cell responses and their implications in IT. More recently, Ji et al. [29] applied trajectory inference, gene regulatory inference, and other computational analyses to 22 baseline and 24 post ICIs treatment samples of stage II/III esophageal squamous cell carcinoma patients to better understand tumor microenvironment changes after therapy. The authors found that upregulated genes in CD8+ effector T cells, which correlated with improved therapy responses, were primarily associated with interferon-gamma (IFNγ) signaling, neutrophil degranulation, and negative regulation of T-cell apoptosis. Additionally, a lower proportion of regulatory T cells (Tregs) was indicative of better treatment response.

This study aims to uncover the transcriptional mechanisms regulating T cell state transitions within the SCC tumor microenvironment, particularly in the context of ICI therapy with PD-1 blockade. Using trajectory inference methods, we analyzed publicly available single-cell RNA-seq data from Yost et al. [30], who previously reported CD8^+^ T clonal replacement and dysfunction. We extended their findings by comprehensively investigating the gene regulatory mechanisms driving all cell state transitions through trajectory inference and regulon inference, with special emphasis on comparing before (PRE-IT) and after (POST-IT) treatment. To address this, we modeled lineage-specific differentiation trajectories for both CD8^+^ and CD4^+^ T cells and reconstructed GRNs that govern T cell dynamics in the TME. Our analysis reveals distinct transcriptional cascades associated with T cell activation and exhaustion and highlights how these networks are rewired following ICI therapy. These findings offer a broader understanding of T cell plasticity and the distinct remodeling dynamics affecting both CD8^+^ and CD4^+^ lymphocyte subsets in response to therapy.

## Results

In this work, we aimed to better understand the molecular mechanisms driving T-cell state transitions upon IT, focusing on the responses to treatment in both CD4^+^ and CD8^+^ T-cell populations. We analyzed a publicly available single-cell transcriptomics dataset (accession number GSE123813), derived from tumors in four patients with squamous cell carcinoma (SCC) treated with ICIs, including data collected before and after treatment from peri-tumoral lymphocytes (CD45^+^ CD3^+^) [30]. Then, we applied state-of-the-art computational tools and a custom analysis pipeline to identify transcriptional regulators of IT-driven cell-state transitions across multiple patients.

A summary of our approach is shown in **Fig. 1**. We began with data exploration and quality control (QC), retaining 25,581 cells that passed quality thresholds for downstream analysis (SI Table S1), followed by unsupervised clustering and cluster annotation to identify cell states based on metadata labels. We then applied trajectory inference to define potential cell-state transitions. We characterized these transitions by identifying variable genes and performing differential gene expression (DEG) analysis. Next, we used the identified DEGs to explore the functional basis of the observed cell-state transitions by performing gene set functional analysis to identify enriched pathways. Finally, we identified transcriptional regulators linked to those trajectories using regulon inference to gain mechanistic insight into the drivers of cell-state transitions.

**Figure 1.**
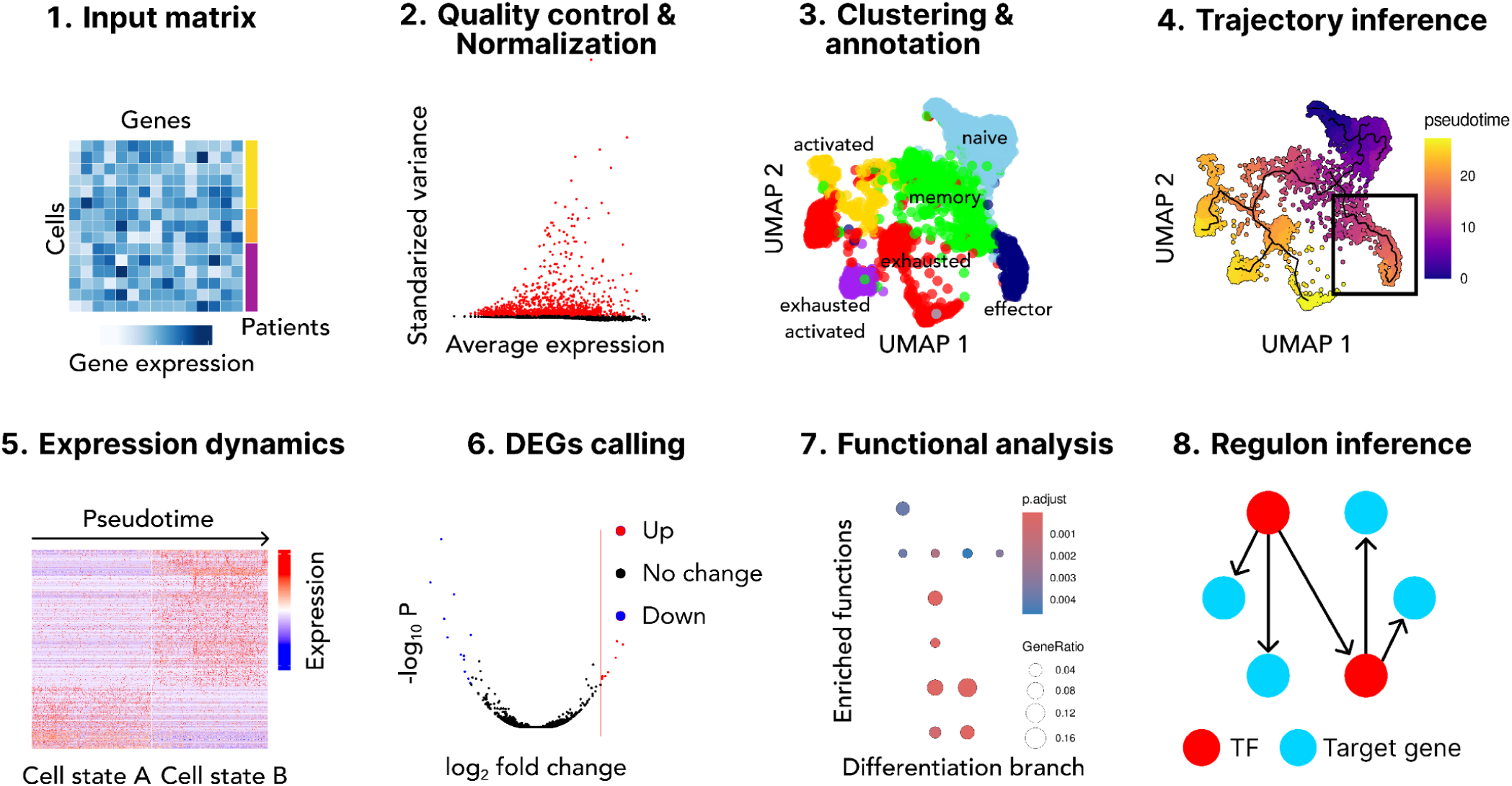
Overall approach used in this study.

### CD8^+^ T cell differentiation trajectories before and after PD-1 blockade therapy

As a first goal, we sought to determine the regulatory processes underlying CD8^+^ T cell differentiation triggered by therapy. We performed quality control and normalization using standard functions (see **Methods** for details) resulting in a filtered dataset of 14,644 CD8^+^ T transcriptome profiles (SI Table S1). We maintained the original annotation based on their canonical marker gene expression defined by Yost et al [30], discriminating CD8^+^ T cells into six subtypes: naive, effector, memory, activated, exhausted and activated/exhausted, populations. We observed no substantial change in the number of CD8^+^ T cells between PRE-IT and POST-IT conditions (**Fig. 2A-B**). Notably, while the naive population accounted for ∼40% of cells in the PRE-IT conditions, this population is virtually absent in the POST-IT conditions. We observed a similar pattern for the effector cells, which shrunk from 11% to only 1%. Another remarkable result is the large increase of memory cells from 13% to 50%, accompanied by an increase in exhausted cells from 23% to 35%. Altogether this observation indicates a deep impact of IT on CD8^+^ T regulation, prompting further investigation into differential regulatory trajectories.

**Figure 2.**
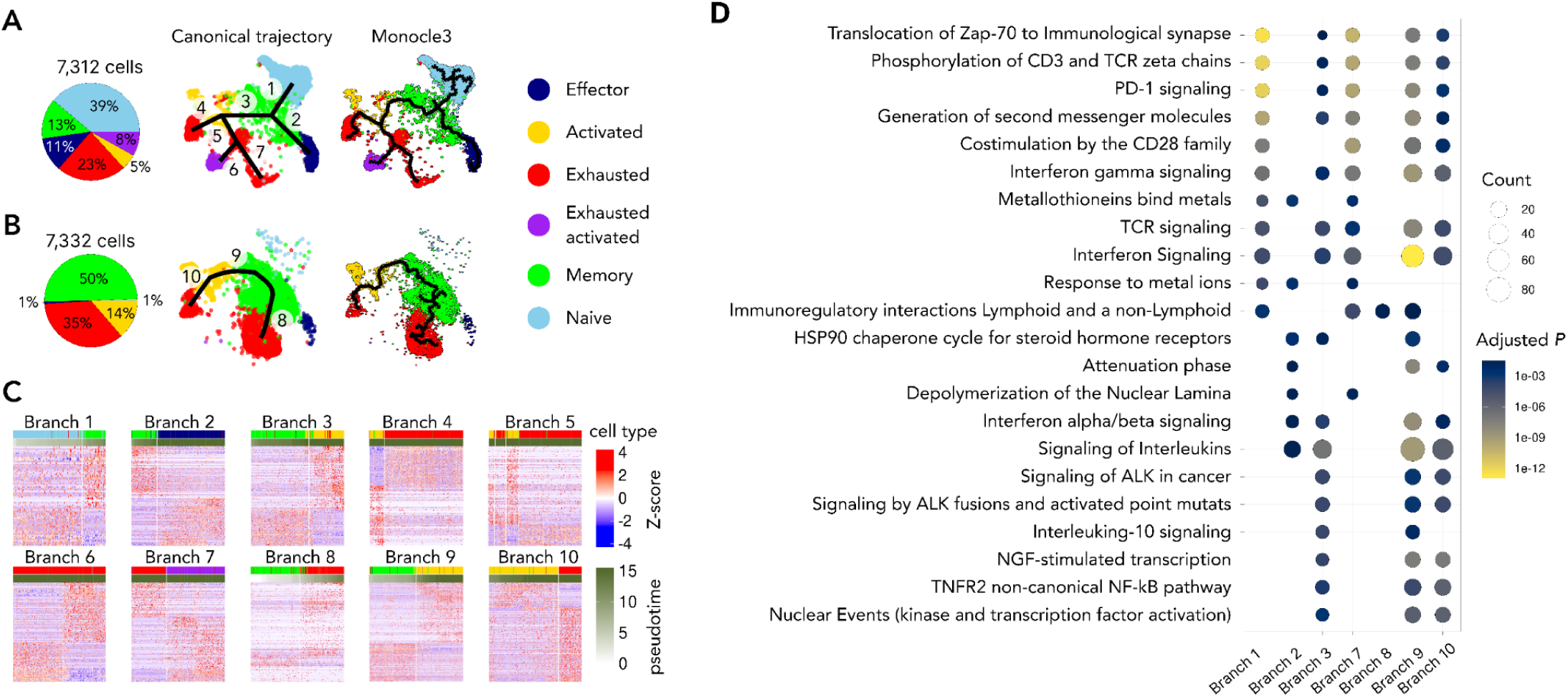
CD8^+^ T cell differentiation trajectories under PRE-IT and POST-IT conditions. **A,B.** Inferred differentiation cell states and trajectories before (A) and after (B) therapy. Pie charts (left panels) indicate cell type proportions. UMAP projections holding expert-curated canonical differentiation trajectories (middle panels; numbers indicate differentiation branches) and Monocle 3-inferred trajectories (right panels). Across all panels, colors indicate cell states. **C.** Expression heatmaps of branch-specific DEGs, with genes (rows) clustered based on expression similarity and cells (columns) ordered by pseudotime progression. **D.** Dot plot visualizing top 10 functional enrichment results for each branch-specific DEG set.

Next, we applied Monocle 3 [20] to infer differentiation trajectories and underlying gene regulatory mechanisms acting on T cell populations within the TME. We used all the cells identified in both PRE-IT and POST-IT and inferred trajectories independently for each dataset. We identified multiple differentiation trajectories labelled numerically in **Fig. 2A-B**. Our analysis revealed distinct trajectory patterns before PRE-IT and POST-IT. For example, while exhausted cells primarily emerged from activated cells in the PRE-IT condition, after therapy they arose not only from activated but also more frequently from memory T cells, highlighting a treatment-induced shift in differentiation dynamics.

To gain a deeper understanding of the transcriptional programs driving differentiation trajectories and cell state transitions, we identified the DEGs (SI Table S2; **Fig. 2C**) and their enriched biological functions (full list of pathways available in SI Table S3; top representative pathways shown in **Fig. 2D**) for each branch across PRE-IT and POST-IT conditions (see **Methods** for details).

Thus, we identified several key genes associated with each differentiation branch. For the naive-to-memory transition prior to therapy (Branch 1), we found 244 DEGs. Notably, we observed enhanced CD28 co-stimulation and TCR signaling which are established hallmarks of early CD8^+^ T-cell differentiation [31]. Altogether, this branch represents the canonical transition from naive to memory T cells and comprises 1,557 cells. In contrast, after therapy, no such transition was observed, suggesting that most cells had already differentiated into a memory state by this point.

It is well established that cytotoxic function within the TME is driven by effector CD8^+^ T cells. Moreover, a subset of effector T cells can transition into long-lived memory cells through dedifferentiation, despite initially acquiring cytotoxic functions [32]. During the transition from memory to effector cells (Branch 2 in **Fig. 2A** and SI Table S2), we identified 1,001 cells and 350 DEGs. The effector cells are characterized by high expression of the surface marker *KLRG1* [33], which was present in our trajectory-specific DEG set, along with key genes such as *CCL3*, *CCL3L3*, *CXCL16*, *CXCR3*, *CXCR4*, *IL10RA*, and *IL9R*. The pathways associated with these cytokines and chemokines support T-cell effector differentiation and regulation [34].

In the transition from memory to activated T cells (Branches 3 and 9 in PRE-IT and POST-IT with 360 and 1,252 DEGs, respectively), we identified the upregulation of *CD69*, one of the earliest cell surface antigens induced upon activation [35]. Additionally, we found a concomitant downregulation of *IL7R*, a crucial marker for memory precursor cells prior to activation [36]. Regarding differentially active pathways in this branch, the *TNFR1-induced NF-kappa-B signaling pathway* (*P* < 2 × 10^−2^) and *Interleukin-12 family signaling* (*P* < 3 × 10^−2^) are specific to PRE-IT and POST-IT conditions, respectively. While the first pathway has been implicated in apoptosis [37], the POST-IT-specific pathway is associated with enhanced cytotoxic activity of CD8^+^ T cells, reflecting therapy-induced augmentation of T cell function [38].

In the transition from activated to exhausted states, we identified Branch 10 under POST-IT conditions, with 711 DEGs and 82 significantly enriched pathways, 12 of which are related to interleukin signaling (SI Table S3). This suggests that cytokines contribute to immune activation and differentiation after treatment. Moreover, we found a significant enrichment for the pathway term *NIK-->noncanonical NF-kB signaling* (*P* < 1 × 10^−2^) in the DEG set for this branch, a mechanism previously reported to sustain the functional fitness of exhausted CD8^+^ T cells [39].

In contrast to what occurs under the POST-IT condition, the transition from activated to exhausted states reveals two distinct branches (Branches 4 and 5) in the PRE-IT condition (**Fig. 2A**). These branches display relatively few DEGs (121 and 109, respectively). However, together with Branch 10, each of all these three branches contain well-known biomarkers of the exhausted phenotype, including the downregulation of *GZMK* and the upregulation of *HAVCR2* [36]. Beyond this complexity, we observed two additional transitions from exhausted cells unique to the PRE-IT state. The first PRE-IT-specific transition, Branch 6, contains only 100 DEGs and leads to a cell state resembling the “exhausted activated” phenotype described by Yost et al. [30]. The second transition branch, Branch 7, involves a subset of exhausted cells giving rise to another exhausted cells subset. This branch includes 385 DEGs and 15 significantly enriched pathways, notably *TCR signaling* (*P* < 3 × 10^−3^), *PD-1 signaling* (*P* < 5 × 10^−10^) and *co-stimulation by the CD28 family* (*P* < 2 × 10^−9^). These findings suggest the existence of two distinct exhausted cell states with differential functional fitness.

Furthermore, unique to the POST-IT condition, we identified a distinct transition from memory to exhausted T cells, corresponding to Branch 8, which contains 441 DEGs. This transition shows the canonical marker shift, including the previously described differential expression of *GZMK* and *HAVCR2*, along with additional exhaustion-associated markers such as *ENTPD1* (*CD39*) and *CTLA4* [40], which are specifically enriched in this branch. Notably, the pathway term *Immunoregulatory interactions between a Lymphoid and a non-Lymphoid cell* is significantly enriched (*P* < 3 × 10^−2^), highlighting the influence of the TME on this differentiation branch. This is consistent with the notion that persistent antigen stimulation is a defining feature of T cell exhaustion within the TME during immunotherapy [41].

This comprehensive analysis highlights not only the complexity of CD8+ T cell dynamics in response to IT, but also the signaling pathways underlying such cell state transitions.

### CD8^+^ T-cell regulatory response before and after therapy

Cell state transitions are driven by changes in gene regulation, with TFs serving as key mechanistic regulators of such shifts. Therefore, we sought to identify the GRNs underlying the previously characterized phenotypic differentiation patterns in PRE-IT and POST-IT conditions in CD8^+^ T cells. To this end, we applied the GRN inference algorithm pySCENIC [42] to the CD8^+^ T-cell single-cell transcriptome profiles described above (see **Methods** for details). In total, we inferred 328 regulons across the entire CD8^+^ T-cell population (**Fig. 3A**; SI Table S4).

**Figure 3.**
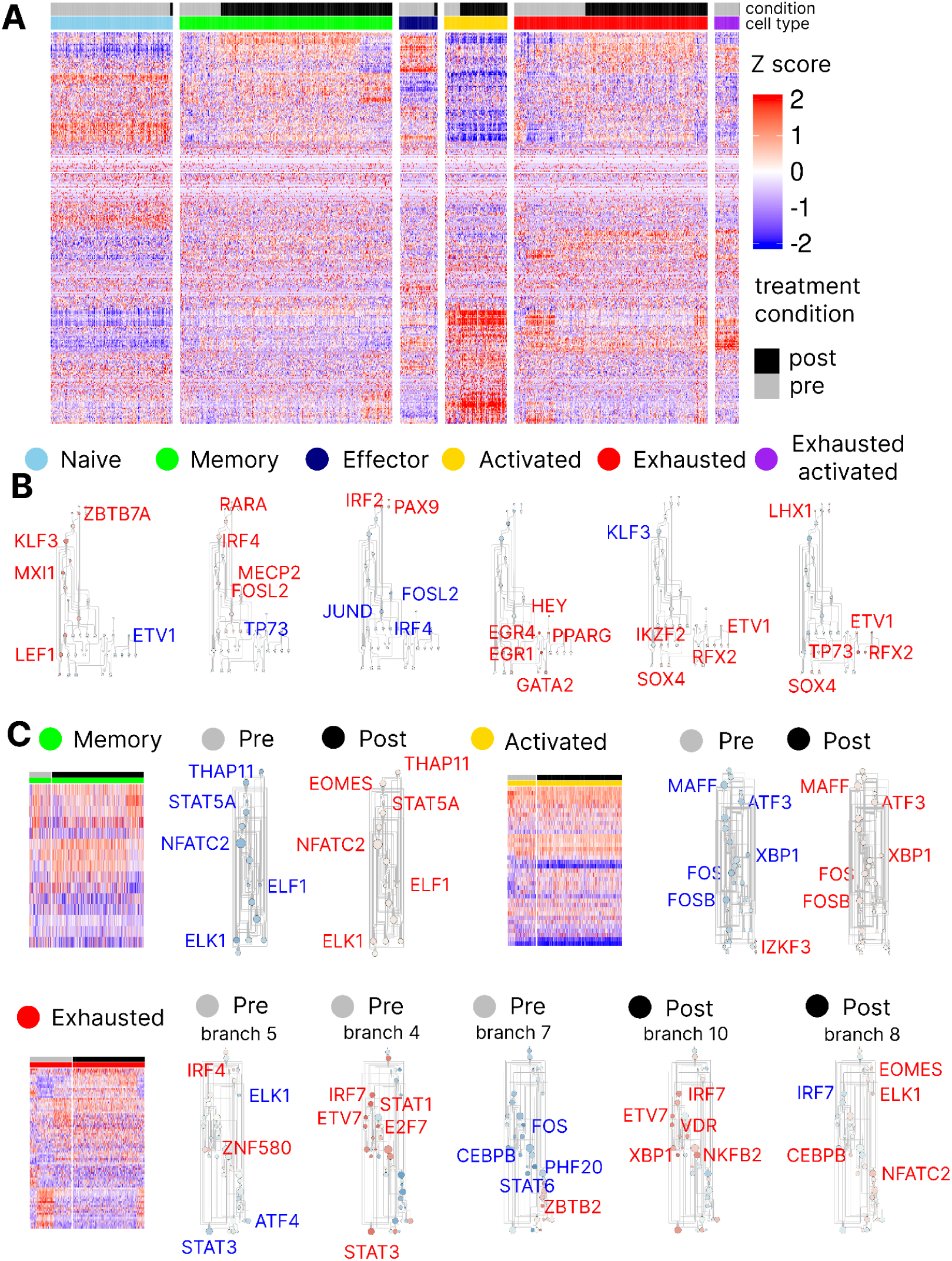
Regulatory dynamics of SCC CD8^+^ T cells before and after therapy. **A.** Heatmap showing relative regulon activity (z-scores) for all 328 regulons inferred by pySCENIC. Top color bars indicate cell type and treatment conditions. The heatmap is partitioned by cell type for visibility purposes. **B.** Cell type-specific TF-TF interaction networks containing the union of the top 10 most specific regulons per cell type. Node color indicates regulon activity (z-score) within cell type, and node size is proportional to interaction degree. Top most differentially active regulons are labelled. **C.** Heatmap of scaled regulon activity for differentially active regulons between PRE-IT and POST-IT conditions, separated by cell type. Adjacent to each heatmap, we show TF-TF interaction networks built as in panel B but for DARs between PRE-IT and POST-IT conditions.

Next, we selected the top 10 most specific regulons for each cell type (<u>SI Table S5</u>). Using these regulons, we constructed a TF-TF interaction network to determine the master regulators that drive differences across CD8^+^ T-cell types (**Fig. 3B**). Among them, we identified KLF2, FOSL2, and BACH2 as key specific regulators in memory cells, all previously shown to promote proliferation and maintenance of the memory phenotype [43,44]. Additionally, we found EGR1 and IKZF2 as key regulators of activated and exhausted cell types, respectively, consistent with previous studies [45,46]. Overall, inferred regulons align well with current knowledge of cell identity.

After characterizing the phenotypic regulatory mechanisms in each cell type, we sought to identify how each CD8^+^ T-cell subtype responded to treatment. To this end, we detected 151 regulons with significantly different activity between treatment conditions (activity difference z-score > 0.5 and Benjamini–Hochberg adjusted *P* < 0.05; see **Methods** for details). Based on these differentially active regulons (DARs), we constructed a TF-TF interaction network specific to each transition (see **Methods** for details). Along the memory cell lineage, we identified 15 DARs (**Fig. 3C**; SI Table S6). Importantly, we observed a significant increase in EOMES activity under POST-IT conditions (*P* < 4 × 10^−35^). EOMES functions maintain memory T-cell identity [47] and cooperates with other TFs as T-bet to induce expression of IL-2 and IL-15, which promote proliferation and persistance in memory T cells [48]. Additionally, we found an increase in STAT5A regulon activity (*P* < 6 × 10^−3^), consistent with its established role in promoting memory CD8^+^ T-cell survival [49]. In activated T cells, we identified 37 DARs (**Fig. 3C**). While the AP-1 regulon was downregulated in PRE-IT conditions, we observed a significant increase in JUN (*P* < 3 × 10^−2^) and FOS (*P* < 2 × 10^−9^) regulon activities under POST-IT conditions. These results indicate enhanced functional state of activated T cells following therapy [50].

Next, we sought to investigate the therapy effects on exhausted cells. Unlike other CD8^+^ T cells, which form a relatively uniform grouping, exhausted cells segregated into multiple distinct clusters in both PRE-IT and POST-IT conditions—three and two clusters, respectively (**Fig. 2A-B**). To quantify the effect of therapy in the GRNs of exhausted CD8^+^ T cells, we adopted a targeted analytical approach. First, we compiled the union of DARs across all five exhausted cell clusters. Then, we identified the significant DARs within each cluster trajectory (130 in total; SI Table S6). Our pseudotime analysis revealed a major shift in the emergence of exhausted T cells in POST-IT compared to PRE-IT conditions. Specifically, only under POST-IT did we identify a cluster of exhausted cells originating from memory cells. This cluster, emerging out of Branch 8, exhibited a significant increase in both TCF7 and EOMES regulon activity (*P* < 4 × 10^−3^ and *P* < 3 × 10^−62^, respectively). These transcription factors are key regulators of Tpex-cell identity, a state associated with improved outcomes in the context of PD-1 therapy [12]. Additionally in POST-IT, we identified a second cluster of exhausted T cells emerging from activated T cells (Branch 10). However, we interpret this cluster as functionally compromised, as it holds significantly reduced (*P* < 2 × 10^−11^) NFATC2 regulon activity—a factor whose absence is linked to T-cell dysfunction [51]. By contrast, PRE-IT conditions displayed a markedly different pattern, with three distinct exhausted T-cell clusters. Among them, the cluster emerging from Branch 7 showed a strong correlation with the most functional competent cluster in POST-IT (PCC = 0.86, *P* < 2 × 10^−27^; **SI Fig. 1**), and retained TCF7 regulon activity. However, the absence of STAT5A and EOMES activity in this cluster emerging from Branch 7 indicates a terminally exhausted phenotype (Tex) [52]. The remaining two exhausted T-cell clusters in PRE-IT, originating from Branches 4 and 5 also displayed signatures of dysfunction. Both clusters showed significantly elevated PRDM1 and IRF4 (*P* < 2 × 10^−2^ and *P* < 1 × 10^−7^, respectively) regulon activity, which are canonical markers of T-cell exhaustion and dysfunction [53]. In particular, high PRDM1 activity has been associated with poor therapeutic efficacy in immune response [54].

Overall, these results highlight the regulatory mechanisms underlying therapy-induced shifts in CD8^+^ T-cell differentiation trajectories and support the emergence of a transcriptionally and functionally enhanced population consistent with Tpex-cell identity.

### CD4^+^ T cell differentiation trajectories before and after therapy

CD8^+^ T cells are the major effectors in IT cancer treatment, yet CD4^+^ T cells play an important role in sustaining adaptive immune responses. Given their critical contribution to anti-tumor immunity, we extended our analysis to CD4^+^ T cells to obtain a more comprehensive view of the mechanisms underlying IT response. Specifically, we used Yost et al. [30] annotations on 10,937 CD4^+^ T cells derived from the same four patients that assemble into four subtypes: naive, T helper type 17 (Th17), T follicular helper (Tfh) and regulatory (Treg). We then followed a similar approach to that applied to CD8^+^ T cells, using Monocle 3 [20] to infer pseudotime trajectories and pySCENIC [42] to identify GRNs (see **Methods** for details).

Cell plasticity is a hallmark of CD4^+^ T cells [55,56], and our analysis provides a clear illustration of this phenomenon. Under PRE-IT conditions, we observed a distinct multifurcation pattern, as naive cells differentiate into three separate arms leading to Th17, Tfh and Treg subsets (**Fig. 4A**). Upon treatment, however, the naive-to-Th17 transition is virtually absent, with this subset decreasing from 21% to only 6% of the total CD4^+^ population (**Fig. 4B**). Interestingly, therapy inverted the relative balance between the two differentiation arms: in PRE-IT, Tregs predominated over Tfh cells (37% vs. 17%; Branches 1 and 2), whereas in POST-IT, Tfh cells became dominant (45% vs. 17%; Branches 4 and 5).

**Figure 4.**
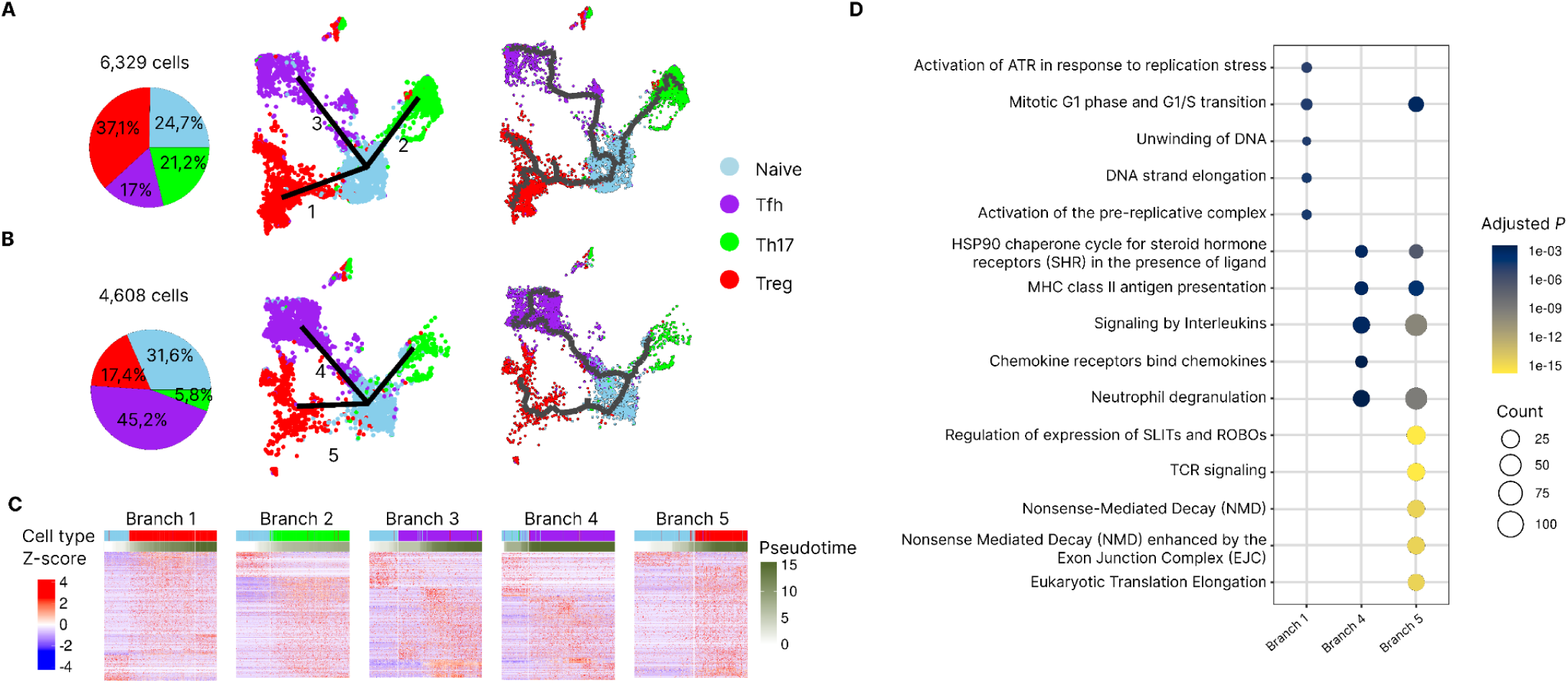
CD4^+^ T-cell differentiation trajectories before and after therapy. **A,B.** Inferred differentiation cell states and trajectories before (A) and after (B) therapy. Pie charts (left panels) show cell-type proportions. UMAP projections display expert-curated canonical differentiation trajectories (middle panels; numbers denote differentiation branches and transitions) and Monocle 3-inferred trajectories (right panels). Across all panels, colors indicate cell states. **C.** Expression heatmaps for branch-specific DEGs. Rows (genes) are clustered by expression, and columns (cells) are sorted by pseudotime. **D.** Dot plot visualizing functional enrichment results for branch-specific DEG sets.

To further characterize differentiation branches in CD4^+^ T cells, we performed differential expression analysis and pathway enrichment (see **Methods** for details). We identified 2,491 DEGs across all branches (**Fig. 4C** and SI Table S7) and 767 significantly enriched pathways (**Fig. 4D** and SI Table S8), representing a diverse set of potential mechanisms driving these transcriptional changes. Quantitatively, POST-IT trajectories involve a markedly larger number of DEGs: while in PRE-IT all branches contained fewer than 200 DEGs each, POST-IT branches exhibited circa 1,200 DEGs each. This pattern likely reflects the pronounced impact of immunotherapy on CD4^+^ T-cell differentiation states and dynamics.

Starting with the trajectories leading to Treg cells, (Branch 1 in PRE-IT and Branch 5 in POST-IT), we identified *FOXP3* upregulation, a well-established master regulator of Treg differentiation [57]. Tregs are classically known for their immunosuppressive and self-regulatory functions [58]. In POST-IT conditions, they appear transcriptionally more active, showing enrichment of key activation pathways such as *Interleukin-2 signaling* (*P* < 1 × 10^−2^) and *TCR signaling* (*P* < 1 × 10^−14^). This increased activation state could help explain the reduced Treg proportions observed in POST-IT. The observation of fewer but more active Tregs in the TME after immunotherapy is biologically nuanced—tumors may adapt to immune pressure by favoring a smaller Treg population in the TME, yet a more potent Treg population, potentially constraining antitumor immune responses [59].

On both branches showing the transition from naive to Tfh cells (Branch 3 in PRE-IT and Branch 4 in POST-IT), we identified *SOX4* as a DEG. Indeed, *SOX4* has been reported to promote inflammation through the differentiation of CXCL13-producing Tfh cells and the formation of tertiary lymphoid structures, which are associated with favorable clinical outcomes [60]. Notably for this transition, we observed *CXCL13* upregulation exclusively under POST-IT conditions. While no pathway term is significantly enriched in PRE-IT (only 158 DEGs were identified in Branch 3), the POST-IT trajectory toward Tfh cells shows significant activation of the interleukin-21 signaling pathway (*P* < 2 × 10^−2^). Together, the increased expression of *CXCL13* and IL21–related signaling indicates a more activated and coordinated immune response, potentially enhancing CD8^+^ T-cell and B-cell recruitment into the TME [61,62].

In Branch 2, which represents the transition from naive cells to Th17 cells, we observed the expected *CD27* downregulation—a molecule known to influence Th17 differentiation and function [63]. This transition, however, could not be reconstructed under POST-IT conditions, likely reflecting the low abundance of Th17-labeled cells (<6%).

Altogether, these results suggest that immunotherapy biases CD4^+^ T-cell differentiation away from the Th17 lineage commitment and toward more transcriptionally active Tregs and CXCL13^+^ Tfh cells, potentially influencing the tumor microenvironment by enhancing immune coordination while maintaining selective immunosuppression.

### CD4^+^ T cell regulatory response before and after therapy

Cell state trajectories in CD4^+^ T cells are governed by dynamic gene-regulatory shifts, primarily mediated by transcription factors. To uncover the regulatory programs driving the distinct differentiation trajectories observed in PRE-IT and POST-IT conditions, we applied the gene regulatory network inference algorithm pySCENIC [42] to the single-cell transcriptomic profiles of CD4^+^ T cells (see **Methods** for details). This analysis identified a total of 331 regulons spanning the entire CD4^+^ T cell population, thereby providing a regulatory map underlying the observed phenotypic transitions (**Fig. 5A**; SI Table S9).

**Figure 5.**
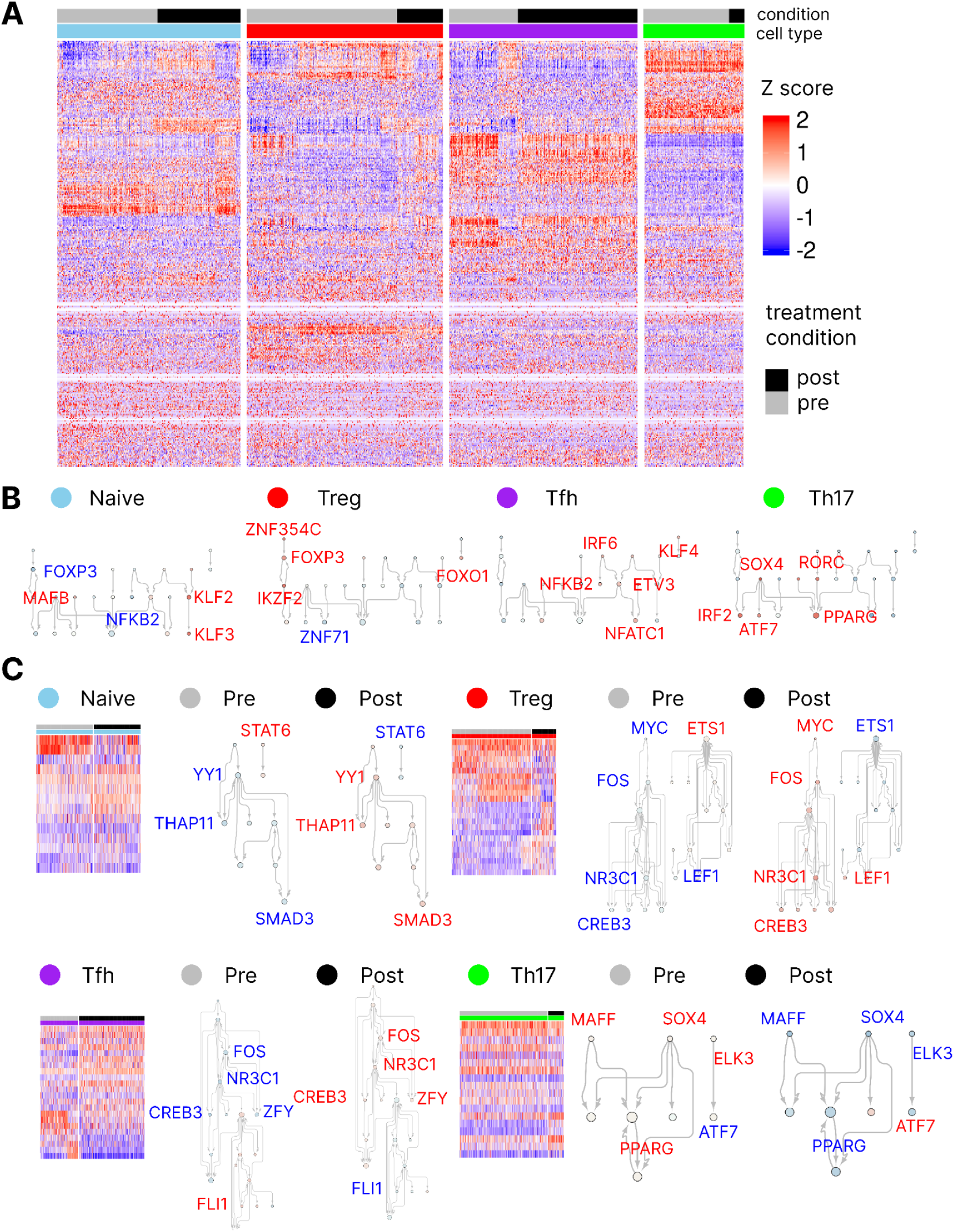
Regulatory landscape of CD4^+^ T cells in SCC before and after therapy. **A.** Heatmap showing scaled regulon activity for all 331 regulons identified by pySCENIC. Top color bars denote cell type and treatment condition. For clarity, the heatmap is split by cell type. **B.** Cell type-specific TF-TF interaction networks constructed from the top 10 most specific regulons per cell subtype. Node color represents regulon activity (z-score), and node size reflects the number of interactions. The top most differentially active regulons are labeled. **C.** Heatmap of DARs between PRE-IT and POST-IT organized by cell type. Next to each heatmap, we show TF-TF interaction networks on differential regulons between PRE-IT and POST-IT conditions.

To deconstruct the regulatory landscape underlying CD4^+^ T cell diversity, we employed the same strategy as previously applied to CD8^+^ T cells. First, we identified the top 10 most cell type-specific regulons and used them to construct a transcription factor–centered regulatory network. This approach revealed a core set of regulators orchestrating distinct CD4^+^ T-cell states (**Fig. 5B**; SI Table S10).

In naive cells, we identified the KLF2 regulon as highly active (activity z-score = 1.24). KLF2 functions as a naive CD4^+^ T-cell–specific transcriptional marker, that maintains quiescence and tissue-homing capacity, and its expression largely disappears following activation [64]. In Tregs, FOXP3 ranked among the most active regulons (activity z-score = 0.93), consistent with its role as the lineage-defining TF for Treg cells [57]. Similarly, in Th17 cells we identified RORC as a highly active regulon (activity z-score = 1.25), in agreement with its well-established role in Th17 differentiation [65]. However, although BCL6 is a known master regulator for Tfh differentiation [66], no regulon could be defined for this TF in the dataset, as its expression was not detected. Nonetheless, NFATC1, which promotes Tfh cell generation [67], is one of the top active regulons in this cell cluster (activity z-score = 0.70). Overall, the identified regulon activity patterns align well with established knowledge from the literature.

Building on our characterization of the regulatory programs defining individual CD4^+^ T-cell phenotypes, we next investigated how these distinct subtypes responded to immunotherapy. We identified 63 DARs (SI Table S11; see **Methods** for details), which we used to construct a TF-TF interaction network.

Along the naive CD4^+^ T-cell lineage, we observed increased YY1 regulon activity in POST-IT conditions. YY1 is known to repress FOXP3 and thereby inhibit Treg differentiation [68]. In addition, observed decreased SMAD3 regulon activity is associated with reduced Th17 differentiation [69], which may partially explain the lower frequency of Tregs and Th17 cells observed after treatment. Under POST-IT conditions, we also detected elevated ETS1 regulon activity, previously described as a positive regulator of Tfh differentiation [66]. The concurrent activation of IRF7 and IRF9 indicates interferon-driven stimulation of Tfh cells, likely reflecting an immunotherapy-induced inflammatory response.

Within the Treg context, the POST-IT regulatory network activity is markedly distinct from that of PRE-IT conditions (**Fig. 5C**). POST-IT samples showed elevated FOXO1 regulon activity, FOXO1 being a TF known to promote Treg survival under nutrient-limited or stress conditions [70]. Together with the concurrent upregulation of FOXP3, these findings indicate that IT profoundly reconfigures Treg regulatory programs.

Within the Th17 cell cluster, we identified SOX4 regulon as significantly downregulated (*P* < 2 × 10^−15^), which is typically associated with naive-like states. This finding suggests the persistence of a transitional population bridging naive and Th17 phenotypes.

Altogether, our findings indicate a shift rebalancing CD4^+^ T-cell states following immunotherapy. The concurrent remodeling of Treg and Tfh populations illustrates how treatment not only promotes effector programs but also modulates regulatory circuits, adding further complexity to the immune landscape of the TME.

## Discussion

While the transcriptional mechanisms driving T cell dynamics in the TME remain incompletely defined, our single-cell and trajectory analyses in SCC data revealed how anti-PD-1 therapy fundamentally remodels both CD8^+^ and CD4^+^ T-cell lineages, uncovering transcriptional regulatory programs underlying treatment response.

Our results show that in PRE-IT conditions, CD8^+^ T cells followed a canonical differentiation route, with naive cells giving rise to both cytotoxic effector and dysfunctional exhausted subsets. In contrast, in POST-IT conditions, we observed a reprogrammed trajectory such that activated T cells upregulated IL-12-associated signaling to enhance cytotoxic responses, and exhausted cells arose from a memory-like progenitor population rather than directly from activated cells. This shift not only suggests a functional restoration of exhausted cells, consistent with recruitment of peripheral T cells into the TME, but also underscores the importance of memory-derived precursors given their superior self-renewal and differentiation capacity [30]. Regulon activity analysis further supported these findings. In POST-IT samples, the Tpex subset emerging from memory cells is characterized by high EOMES and TCF7 activity, both linked to better clinical outcomes [28]. Conversely, terminally exhausted clusters showed reduced JUN, FOS, NFAT and increased PRDM1 regulon activity, consistent with irreversible dysfunction. Our results uncovered further additional TF candidates that may also drive this differentiation trajectory.

In PRE-IT conditions, naive CD4^+^ T cells exhibited the expected classic multifurcation into Th17, Tfh, and Treg lineages, reflecting their well-known plasticity [55,56]. However, POST-IT conditions lacked a clear Th17 branch (not found by Monocle 3), while Tfh and Treg proportions were inverted, indicating a likely therapy-induced rewiring of differentiation trajectories. Mechanistically, POST-IT naive cells displayed increased YY1 and reduced SMAD3 regulon activity, consistent with inhibited Treg and Th17 differentiation [68,69]. Although Treg frequencies declined, their remaining population became transcriptionally more active, showing increased FOXP3 and FOXO1 activity and enrichment of TCR and IL-2 signaling pathways, suggesting enhanced survival and suppressive potential within the TME. Meanwhile, Tfh cells activated ETS1, IRF7, and IRF9 regulons and upregulated CXCL13 and IL-21 pathways, consistent with an interferon-driven effector program that may facilitate CD8^+^ T-cell recruitment for coordination of antitumor immunity.

While our study leverages the same SCC single-cell RNA-seq dataset reported by Yost et al. [30], our analytical perspective is different. Yost et al. focused primarily on TCR repertoire dynamics, revealing clonal replacement of tumor-specific CD8^+^ T cells following PD-1 blockade in basal cell carcinoma. In contrast, we centered our approach on transcriptional state transitions, integrating trajectory inference and gene regulatory network modeling across both CD8^+^ and CD4^+^ T cells. This systems-level approach enabled a more comprehensive reconstruction of the differentiation dynamics and the transcriptional programs that govern them. As a result, we identified key regulatory factors and signaling pathways shaping post-therapy T cell states, providing mechanistic insights into treatment-induced immune remodeling that were beyond the scope of the original study.

As expected, we identified the TFs that drive each cell type transition, with the exception of BCL6 in the Tfh lineage. This limitation likely arises because mechanistic inference GRN methods such as pySCENIC rely on TF-target gene binding evidence that is cell-type context specific. Integrating additional regulatory layers, such as cell-type resolved ATAC-seq data, could help refine these predictions. However, even multi-omics GRN inference approaches remain limited in their ability to infer direct mechanistic interactions, and experimental validation methods such as ChIP-seq are ultimately required to confirm these predicted regulatory links [71].

Another limitation of our study is that available single-cell profiles were obtained from only four patients, which may not fully capture the inter-individual heterogeneity in treatment responses. Also, we analyzed over 13,000 CD8^+^ and 9,000 CD4^+^ T cells, but rarer subsets such as Th17 cells remained underpowered, potentially reducing trajectory resolution and differential expression sensitivity. Moreover, although overall cell yields were comparable between PRE-IT and POST-IT conditions, some individual patients exhibited markedly reduced cell recovery in POST-IT. Therefore, larger and more diverse patient cohorts will be essential to validate and generalize observed trajectory and regulon patterns across tumor types and patient populations [72].

Furthermore, pseudotime estimation and branch assignment inherently depend on the assigned cell-type annotation. In addition, as implemented in Monocle 3, inferred trajectories rely on graph structures embedded in the non-linear dimensionality reduction methods, such as UMAP [73]. Although manual curation and established biomarkers guided our branches labeling, potential misclassifications or oversimplified subtype definitions could bias the inferred differentiation paths. Future improvements may come from standardized annotation frameworks and multimodal trajectory algorithms that integrate complementary molecular layers to enhance robustness [74].

In summary, PD-1 blockade fundamentally reprograms T-cell immunity within the TME, establishing a distinct memory-to-Tpex differentiation route in CD8^+^ T cells, driven by increased EOMES and TCF7 activity, while concurrently redirecting CD4^+^ subsets away from Th17 and Treg lineages. Despite their reduced abundance, Tregs in POST-IT conditions exhibit heightened transcriptional activity, and naive CD4^+^ cells preferentially commit toward CXCL13^+^ Tfh differentiation. Together, this dual remodeling points to a restoration in cytotoxic and proliferative potential in exhausted CD8^+^ cells while enhancing the helper and regulatory phenotypes in CD4^+^ cells, thereby coordinating the cellular and humoral arms of the anti-tumor immune response.

Clinically, tracking the emergence of memory-derived Tpex cells (EOMES^+^/TCF7^+^), the CXCL13^+^ Tfh expansion, and the reduction yet heightened activation of Tregs offers a composite biomarker signature to of therapy effect. Moreover, intervention strategies that enhance the EOMES–TCF7 axis or the IL-21/IFN-driven Tfh program could further amplify effector functions, whereas selective modulation of FOXP3^+^ Treg fitness may help limit immunosuppression. Together, these findings define actionable transcriptional routes that could inform next-generation combination regimens aimed at sustaining long-term tumor control.

Overall, our findings illustrate how PD-1 blockade remodels the T cell landscape within the TME. However, at this point establishing causal links between these molecular changes and clinical outcomes remains challenging. Future studies combining longitudinal single-cell profiling with clinical response data in larger patient cohorts will be essential to validate these mechanisms and refine predictive biomarkers for improved immunotherapy design.

## Methods

### Data accession and quality control

We obtained single-cell RNA-seq expression data from the Gene Expression Omnibus (GEO; accession GSE123813), comprising tumor-infiltrating lymphocytes from four patients, sampled before (PRE-IT) and after (POST-IT) anti-PD-1 therapy. The dataset included author-provided metadata and cell-type annotations based on canonical marker gene expression. Following Seurat (v5.3.0) guidelines, we examined feature count distributions per cell and mitochondrial and ribosomal read fractions to ensure data quality. In line with established quality control standards [75], we retained genes with non-zero expression in at least five cells, retained cells with ≤12,000 total counts and 1,000–4,000 detected genes. Cells with >8% mitochondrial gene content were excluded. Raw counts were then normalized and scaled using Seurat’s default pipeline. Principal component (PC) analysis was performed on the top 2,000 variable genes, retaining the first 16 PCs. Unsupervised clustering was performed at a resolution of 0.3, and UMAP embeddings were generated using the same PCs. All the analyses were conducted in R (v4.4.0) through RStudio.

### Trajectory inference

Following quality control, the filtered Seurat object containing normalized gene counts and metadata was converted into a Monocle 3 `cell_data_set` object. Trajectory preprocessing and graph learning were performed using Monocle 3 (v1.4.26) default settings [10]. Then, pseudotime was initiated (rooted) at the naive T-cell population in all conditions, as this subset represents the earliest stage in T-cell differentiation (in this dataset) according to established literature [76]. In POST-IT CD8^+^ trajectories, however, the memory T-cell cluster was designated as the trajectory origin for trajectory inference calculations due to the low abundance of naive cells.

### Differential expression and functional enrichment analysis

Differential expression analysis was performed for within each trajectory branch by subsetting cells assigned to that branch and comparing dedifferentiated versus differentiated states using Seurat’s Wilcoxon rank-sum test (`FindMarkers`). *P* values were adjusted using the Benjamini–Hochberg correction [77], and genes with |log₂FC| > 0.5 and an adjusted *P* < 0.05 were retained. Significant DEGs for each branch were subjected to pathway enrichment analysis using clusterProfiler (v 4.12.6) [78] on Reactome Pathway ontology [79]. Pathways with Benjamini–Hochberg–adjusted *P* < 0.05 were considered significantly enriched.

### Gene regulatory network inference and TF–TF network construction

Gene regulatory networks were inferred using pySCENIC (v0.12.0) [39] (Docker container) independently for CD8^+^ and CD4^+^ T-cell expression matrices. The analysis followed the standard three-step workflow: (1) co-expression module detection (adjacency matrix inference), (2) cis-regulatory motif pruning to refine TF–target interactions, and (3) regulon activity scoring to quantify TF regulon enrichment. From the resulting regulons, we selected the ten most cell type–specific regulons ranked by Jensen-Shannon divergence, to serve as marker TFs for each cell type.

### Differential regulon activity identification

Cell types containing at least 200 cells in both PRE-IT and POST-IT conditions were retained for differential regulon activity analysis. For each regulon, median activity values were computed per cell type and treatment condition, and those showing an absolute median difference greater than 0.5 were tested using two-sample t-tests. *P* values were adjusted using the Benjamini–Hochberg correction, and regulons with adjusted *P* < 0.05 were considered significantly differentially active.

### TF-TF interaction network visualization

TF-TF interaction networks were visualized in Cytoscape (v3.10.2) [80] by importing the regulon adjacency matrix derived from pySCENIC. Node size was mapped to the TF interactions degree and node color to regulon activity (median z-score for the particular cell subset). We rendered networks using a hierarchical layout implemented in the *yFiles Hierarchic Layout* algorithm (v1.1.5).

## Code availability

All computational tools and scripts used to reproduce results presented in this study are publicly available at the GitHub repository https://github.com/rogercasalsfr/Itx_anl.

## Acknowledgments

The authors gratefully acknowledge Christian Brander and Alex Olvera for their insightful discussions and valuable suggestions during the development of this study.

## Data availability

All data generated or analysed during this study are included in this published article (and its Supplementary Information files).

## Authors contributions

RCF: Conceptualization, data curation, formal analysis, funding acquisition, investigation, methodology, software, validation, visualization, writing – original draft and writing – review & editing.

JVF: Conceptualization, funding acquisition, project administration, resources, supervision, writing – original draft and writing – review & editing.

LN: Conceptualization, funding acquisition, project administration, supervision, writing – original draft and writing – review & editing.

ALGDL: Conceptualization, funding acquisition, project administration, supervision, writing – original draft and writing – review & editing.

